# Bio-On-Magnetic-Beads (BOMB): Open platform for high-throughput nucleic acid extraction and manipulation

**DOI:** 10.1101/414516

**Authors:** Phil Oberacker, Peter Stepper, Donna M Bond, Sven Höhn, Jule Focken, Vivien Meyer, Luca Schelle, Victoria J Sugrue, Gert-Jan Jeunen, Tim Moser, Steven R Hore, Ferdinand von Meyenn, Katharina Hipp, Timothy A Hore, Tomasz P Jurkowski

**Author notes:** Co-first authors (Phil Oberacker, Peter Stepper, Donna M Bond).

## Abstract

Current molecular biology laboratories rely heavily on the purification and manipulation of nucleic acids. Yet, commonly used centrifuge-and column-based protocols require specialised equipment, often use toxic reagents and are not economically scalable or practical to use in a high-throughput manner. Although it has been known for some time that magnetic beads can provide an elegant answer to these issues, the development of open-source protocols based on beads has been limited. In this article, we provide step-by-step instructions for an easy synthesis of functionalised magnetic beads, and detailed protocols for their use in the high-throughput purification of plasmids, genomic DNA and total RNA from different sources, as well as environmental TNA and PCR amplicons. We also provide a bead-based protocol for bisulfite conversion, and size selection of DNA and RNA fragments. Comparison to other methods highlights the capability, versatility and extreme cost-effectiveness of using magnetic beads. These open source protocols and the associated webpage (https://bomb.bio) can serve as a platform for further protocol customisation and community engagement.

## Introduction

The nucleic acids, RNA and DNA, essentially harbour all heritable biological information so far known [1]. As such, the majority of molecular biology laboratories are dependent upon nucleic acid purification and manipulation. Initial purification can be carried out from a variety of sources, which commonly include bacterial cultures, plant and animal cells or tissues. Nucleic acids may also be derived from cell-free sources, such as blood plasma, various environmental substrates, or from *in vitro* reactions like PCR. Once purified, nucleic acids are usually manipulated in some manner ahead of analysis or functional use – a process which in turn requires further appropriate reagents and handling. Most current nucleic acid purification and manipulation techniques rely upon either a commercially produced silica-based column, or a centrifuge (often both). In addition to commonly using toxic chemicals such as phenol, these protocols are generally not suitable for high-throughput approaches, whereby ≥96 samples are processed simultaneously. This is because standard benchtop centrifuges only hold 24 tubes and multi-channel pipettes or liquid handling robots cannot be used to accelerate the isolations. Moreover, the per sample cost of silica columns make processing large numbers prohibitively expensive.

Magnetic beads are small nano-or micro-particles and have long been recognised as a way to solve scalability issues with respect to nucleic acid purification and manipulation [2–4]. Their most useful characteristic is the ability to achieve solid-phase reversible immobilisation (SPRI) [3]; meaning they can reversibly bind nucleic acid under dehydrating conditions and, when in the presence of a strong magnet, can be safely immobilised throughout multiple wash and manipulation steps. Magnetic bead protocols are inherently scalable due to the fact that they are independent of centrifugation and the required materials are exceedingly cheap both to purchase and manufacture in a laboratory setting. However, despite these attractive attributes, surprisingly little community effort has been committed to the development of open-source protocols featuring their use.

Here, we present Bio-On-Magnetic-Beads (BOMB), an open-source platform consisting of both novel and existing magnetic bead-based protocols that are capable of a wide-range of nucleic acid purification and manipulation experiments (Fig 1). We first detail a method for simple synthesis of magnetic nanoparticles and their functionalisation with either a silica-or carboxyl-coating, that can be performed in any molecular biology laboratory with standard equipment. We further show how cheap magnetic immobilisation devices can be assembled or fabricated for 1.5 ml microcentrifuge tubes and 96-well plates. Finally, we provide protocols for use of the beads and magnetic racks in purifying bacterial plasmid DNA, total nucleic acid (TNA), genomic DNA, PCR amplicons, environmental DNA, as well as total RNA from various sources, all of which have been validated in both modular and high-throughput settings. We have also developed an open-source protocol for bisulfite conversion of DNA used in epigenetic analysis and update existing protocols for size selection of DNA fragments.

**Fig 1.**
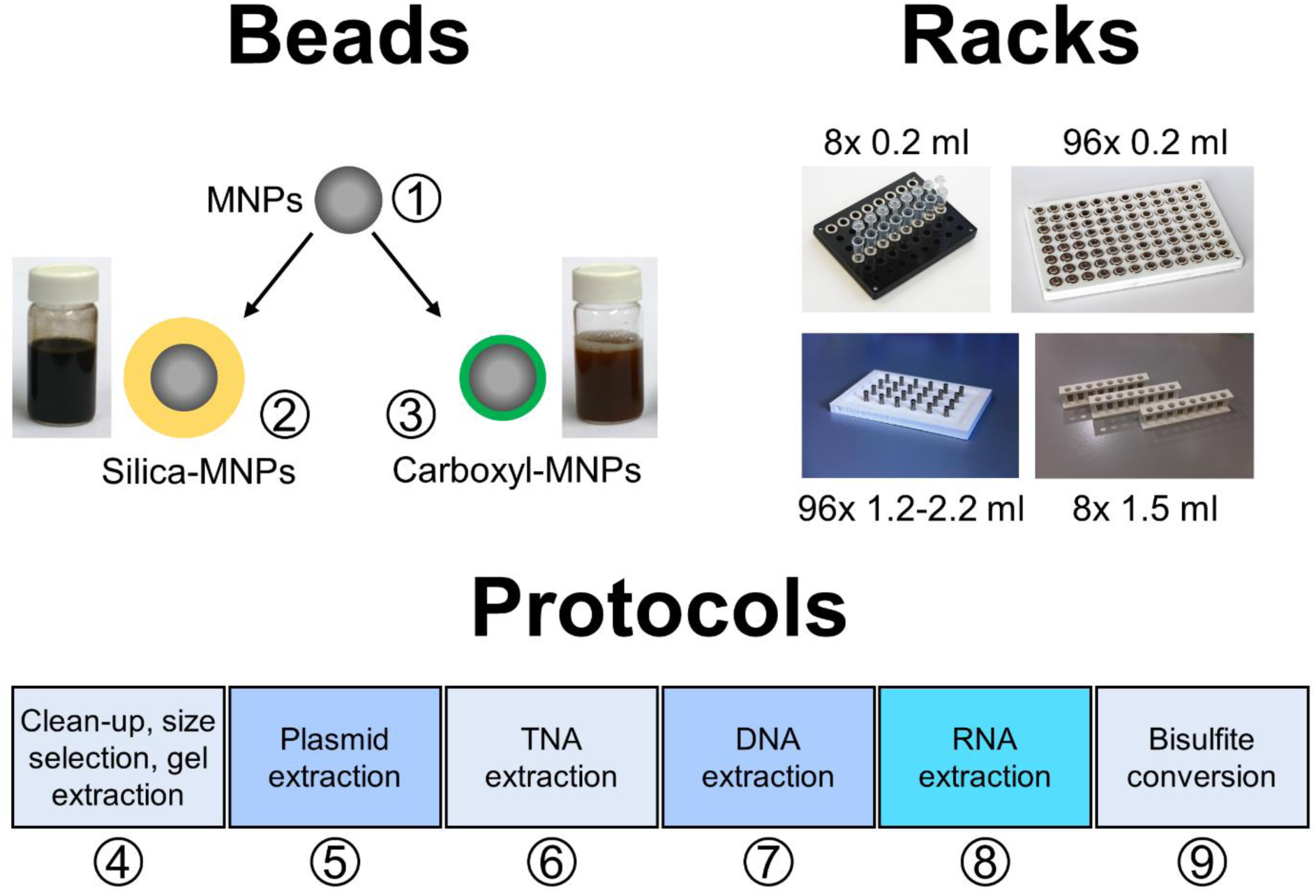
The BOMB platform. The Bio-On-Magnetic-Beads platform is composed of magnetic ferrite nano-particles (MNPs, 1) that can be coated with either a silica (2) or carboxylate surface (3), BOMB magnetic racks produced by laser cutting or 3D printing and the basic BOMB protocols for purification of nucleic acids from various sample origins (4-9). The circled numbers indicate the protocols for the respective procedure.

In the expectation that BOMB protocols will benefit from continued refinement and development, we provide a forum-type website, which allows community engagement (https://bomb.bio). Given the impressive economic advantages of magnetic beads for nucleic acid extraction and manipulation, both in terms of capital outlay and per sample costs, we consider the BOMB platform a positive step towards the democratisation of life sciences.

### Building the BOMB molecular biology platform

The essential components of a magnetic bead platform are the beads themselves and a magnet strong enough to immobilise them. While many life science researchers will be familiar with proprietary beads (e.g. DynaBeads® for antibody capture, AMPure® beads for size selection), few are aware that they can assemble both beads and magnet components themselves from cheap materials. However, in order to do so, some potentially unfamiliar concepts need to be explained.

Firstly, magnetic beads commonly used for molecular biology come in two major forms; either relatively small (50 nm-2 µm) particles constructed from a solid ferrite core, or larger ferrite-polymer combinations (1-5 µm) [5]. Both bead types work well for nucleic acid purification and manipulation, however their different physical and chemical properties do change their behaviour. For example, the polymer within the larger ferrite-polymer beads effectively lowers the density of the bead so they are less likely to settle out of the suspension during handling steps. The smaller solid core ferrite beads have a larger relative surface area for binding and can also be easily made in a standard molecular biology laboratory (see protocols below).

A key aspect of magnetic beads used for molecular biology is that, irrespective of their size, they need to be chemically coated. The first reason for doing this is to provide stability for the bead - without coating, oxidation of the ferrite would lead to contamination of potentially sensitive samples with iron ions and the beads would lose their magnetic properties over time. In addition, chemical coating grants additional function to magnetic beads. For example, silica-or carboxylated-polymer coatings are most commonly used because in addition to providing bead stability, they are relatively chemically inert (silica) or negatively charged (carboxylate), thus facilitating desorption of the negatively charged nucleic acids from the beads during elution steps. Here, we outline a simple protocol for preparation of silica-or carboxylate-coated beads in a standard life science laboratory, and production of magnetic racks suitable for their immobilisation.

### Simple synthesis of functionalised magnetic beads for nucleic acid manipulation Protocol #1: Synthesis of ferrite core magnetic particles

Ferrite nanoparticles can be synthesised using various protocols (reviewed in [6,7]). We adopted the broadly used co-precipitation method due to its simplicity and efficiency, but also because it does not require any specialised equipment [8]. Briefly, FeCl_3_ and FeCl_2_ solutions are mixed in 1:2 molar ratio and slowly dripped into a preheated alkali solution leading to the formation of black ferrite (Fe_3_O_4_) precipitate (Supplementary protocol #1). The ferrite particles synthesised using this approach have a diameter of ∼5-20 nm as judged by transmission electron microscopy (TEM) images (Fig S1 A). Oxygen is known to interfere with the ferrite precipitation reaction, therefore the alkali solution used should be degassed and heated to >80 °C. After synthesis, the core particles are extensively washed with deionised water. In order to prevent oxidation of the ferrite, we recommend coating the beads immediately after synthesis. However, it is also possible to stabilise them in the short term using detergents, sodium oleate, polyvinylpyrrolidone (PVP) or other chemicals (reviewed in [6,7]) and upon lyophilisation can be stored in an air-tight container under inert atmosphere.

### Protocol #2: Coating magnetic nanoparticles with silica

In similar fashion to storing solutions inside glass bottles, encasing ferrite nanoparticles in silica prevents magnetic bead oxidation and leakage of iron ions. Silica-coating also provides an inert surface for precipitation of nucleic acid without the risk of irreversible association. In order to coat ferrite nanoparticles with silica, we use a modified Stöber method [9], in which tetraethyl orthosilicate (TEOS) is hydrolysed in a basic environment, thus forming a SiO_2_ layer surrounding the magnetic core. The thickness of the silica coat (and therefore the size of the particle) can be controlled through the addition of increasing amount of TEOS [10]. The provided standard coating protocol results in the silica-coated beads with a size of approximately 400 nm (Fig S1 B), which perform well for a wide range of the nucleic acid purification and manipulation experiments (Supplementary protocol #2.1).

### Protocol #3: Coating magnetic nanoparticles with methacrylic acid (carboxyl-coating)

An alternative way to stabilise magnetic particles is to coat them with carboxylate modified polymer (Fig S1 C, D). While potentially not providing the same stability as silica, carboxylate-coating endows the ferrite core with a weak negative charge, thus altering its electrostatic interaction with nucleic acids and ultimately affecting bead functionality. Although other reaction schemes are possible, we polymerise methacrylic acid monomers on top of the magnetic nanoparticles, thus providing a negatively charged carboxylated coat. For this, the ferrite core particles are dispersed with a detergent (sodium dodecyl sulfate) and a layer of polymethacrylic acid (PMAA) is deposited on the surface of the beads by a free-radical retrograde precipitation polymerisation reaction [11] (Supplementary protocol #3.1 and #3.2).

### For those in a hurry: buy commercially available beads

Although making your own magnetic beads is by far the most economical way to make use of magnetic beads for nucleic acid handling, for many laboratories, bead-based systems can still be developed relatively cheaply from commercial sources (numerous beads are available, however, where used in protocols here, we purchased Sera-Mag SpeedBead Carboxylate-Modified Magnetic Particles, Hydrophylic, 15 ml, cat., 45152105050250). A single bottle of carboxylate functionalised beads can be purchased for approximately the same cost as a DNA extraction kit based on plastic silica columns for 50-100 samples. However, the purchased beads can provide purification of DNA, RNA or total nucleic acid (TNA) from up to 40,000 samples. While there are advantages and disadvantages to either system, we find that our laboratory-made and commercially sourced beads show similar performance.

### Immobilisation racks for magnetic separation

Magnetic nanoparticles are so useful because they can be immobilised and re-suspended easily by moving them in and out of a magnetic field. The simplest way to do this is using immobilisation racks consisting of magnet arrays, either in microtube or microplate format. Despite their simple construction and lack of moving parts, commercially available immobilisation racks are surprisingly expensive. However, they can be assembled in a laboratory setting relatively easily and cheaply. The most critical component of the rack is the magnet itself. In order to achieve rapid separation, it is best to use high-quality neodymium magnets, N42 grade or above. We have created a number of different racks that are suitable for specific vessel formats using recycled materials or custom 3D-printed/laser-cut parts (Fig 1). For example, when using 8-strip PCR tubes, 96-well microplates and deep-well plates, ring magnets can be fixed onto the top of old pipette tip box components using cyanoacrylate adhesive (e.g. “superglue”), or can be jammed in between wells of the plate. For those who have access to laser cutting or 3-D printing, we have designed 96-well racks that are suitable for a range of deep-well plates and a 3D-printable plastic rack which can hold up to 8 microcentrifuge tubes (Supplementary protocol #A).

### BOMB platform protocols

Once the basic BOMB tools (i.e. beads and magnetic racks) have been created or purchased, there is a broad variety of molecular biology experiments that can be performed with only basic chemical supplies. Many experiments in molecular biology laboratories start with purification of nucleic acid from source cells or tissues (Fig 2) and go on to perform some type of manipulation (often involving many intermediate purification steps) prior to final quantitation or introduction back into a biological system. Magnetic bead-based protocols can mediate each of these purification and manipulation steps in both a universal and modular fashion (Fig 3). This means that for almost any application, reliance upon purchased kits can be dramatically reduced or removed entirely. Here, we highlight some of the most commonly used protocols and their utility. Detailed protocols can be found in the supplementary information.

**Fig 2.**
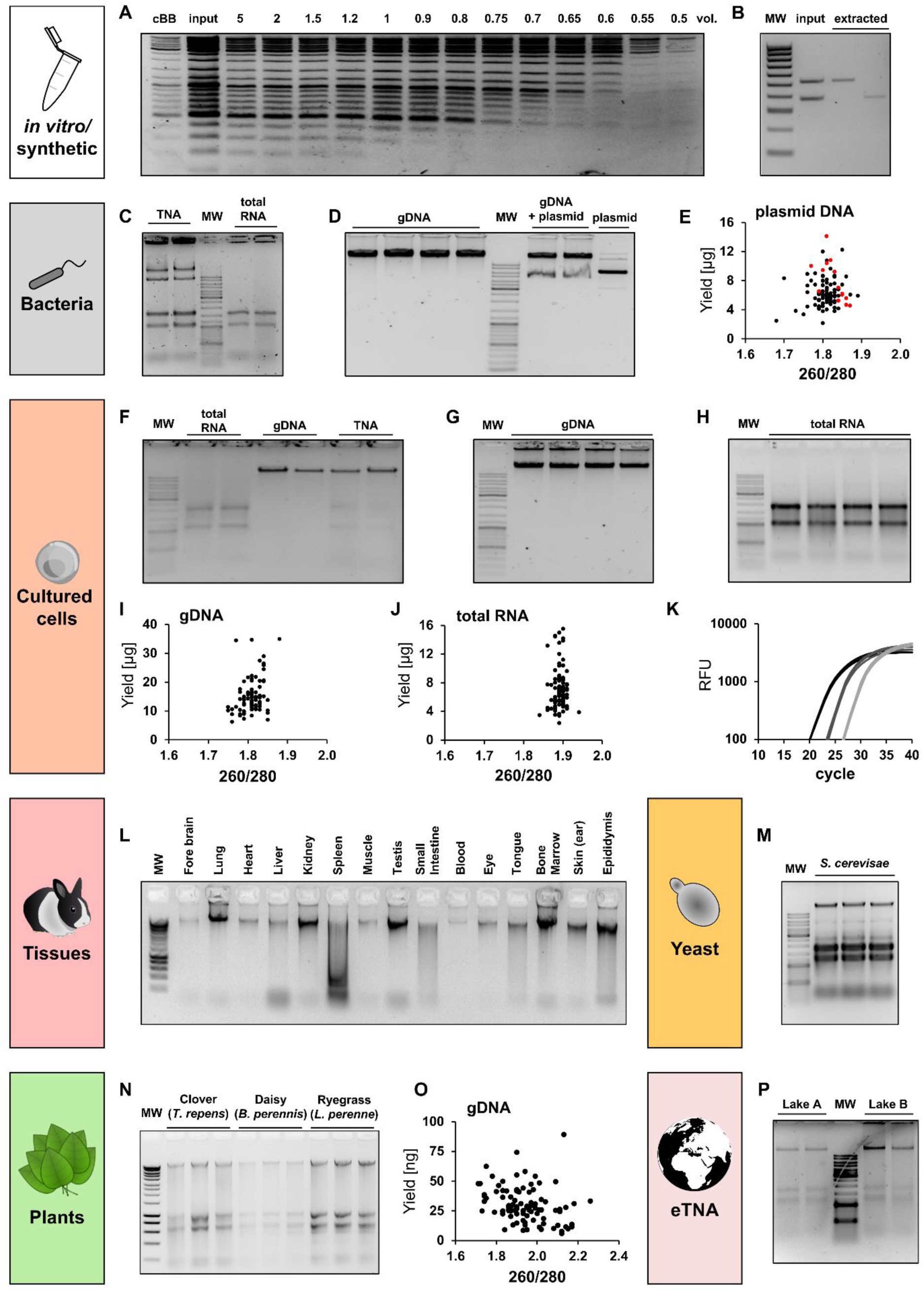
Versatility of the BOMB protocols for nucleic acid isolation. *Nucleic acid extraction from various sample origins using the BOMB extraction protocols (#4-8)*. *(A) Size exclusion of GeneRuler DNA Ladder Mix (Thermo) using BOMB silica-coated magnetic beads (BOMB protocol #4.1). Different volumes of binding buffer compared to sample volume were used to achieve size exclusion. 2 volumes of commercial binding buffer (cBB) were used as a control relative to input. (B) Gel extraction of CHD1 PCR products using female chicken (*Gallus gallus domesticus*) gDNA as a template. The two rightmost lanes contain the gel extracted bands from lane two using BOMB protocol #4.3 with carboxyl-coated magnetic beads. The volumes loaded are proportional (i.e. the right hand 2 lanes represent the efficiency of capture from the left hand lane). MW: Hyperladder IV (Bioline). (C) TNA isolation (BOMB protocol #6.6) from* E. coli *(left two lanes) followed by DNase I digest (right two lanes). MW: GeneRuler DNA Ladder Mix (Thermo). (D) Genomic DNA isolated from TOP10* E. coli. *The first four lanes contain purified DNA from untransformed cells (BOMB protocol #7.1), whereas the three rightmost lanes contain the DNA of two pellets of* E. coli *transformed with a pHAGE-EFS-insert plasmid and also the purified plasmid itself (BOMB protocol #5.1). MW: GeneRuler DNA Ladder Mix (Thermo). (E) Total plasmid DNA yield [µg] extracted from* E. coli, *plotted against the A260 nm/A280 nm ratio for each sample. DNA concentration and purity were measured with UV-Spectroscopy (NanoDrop). Black dots represent samples extracted using the BOMB plasmid extraction protocol #5.1, red dots represent samples processed using a commercial kit. (F) Isolation of total nucleic acid (TNA) from HEK293 cells using the BOMB protocol #6.1 (lanes 6+7) followed by digestion with either DNase I (lanes 2+3) or RNase A (lanes 4+5). (G) Genomic DNA isolated from 500K HEK293 cells using BOMB protocol #7.1. (H) Total RNA isolated from 500K HEK293 cells following BOMB protocol #8.1. (I) Total DNA yield [µg] of representative extractions from HEK293 cells, plotted against the A260 nm/A280 nm ratio for each sample. (J) Total RNA yield [µg] of representative extractions from HEK293 cells, plotted against the A260 nm/A280 nm ratio for each sample. (K) qPCR amplification curve of a 10-fold serial dilution (black: undiluted, dark grey: 1:10 diluted, light grey: 1:100 diluted) of RNA from HEK293 cells reverse-transcribed into cDNA. (L) Genomic DNA isolated from various rabbit (*Oryctolagus cuniculus*) tissues using BOMB silica-coated magnetic beads following BOMB protocol #6.3. MW: Hyperladder I (Bioline). (M) TNA isolation from yeast (*S. cerevisae*) using BOMB protocol #6.5. MW: GeneRuler DNA Ladder Mix (Thermo). (N) TNA isolation from clover (*Trifolium repens*), daisy (*Bellis perennis*) and ryegrass (*Lolium perenne*) according to BOMB protocol #6.4 with carboxyl-coated magnetic beads. MW: Hyperladder I (Bioline). (O) Total DNA yield [ng] of representative extractions from leaves of* T. repens, plotted against the A260 nm/A280 nm ratio of each sample. (P) TNA isolation from 50 ml of lake water following BOMB protocol #6.7. MW: GeneRuler DNA Ladder Mix (Thermo).

**Fig 3.**
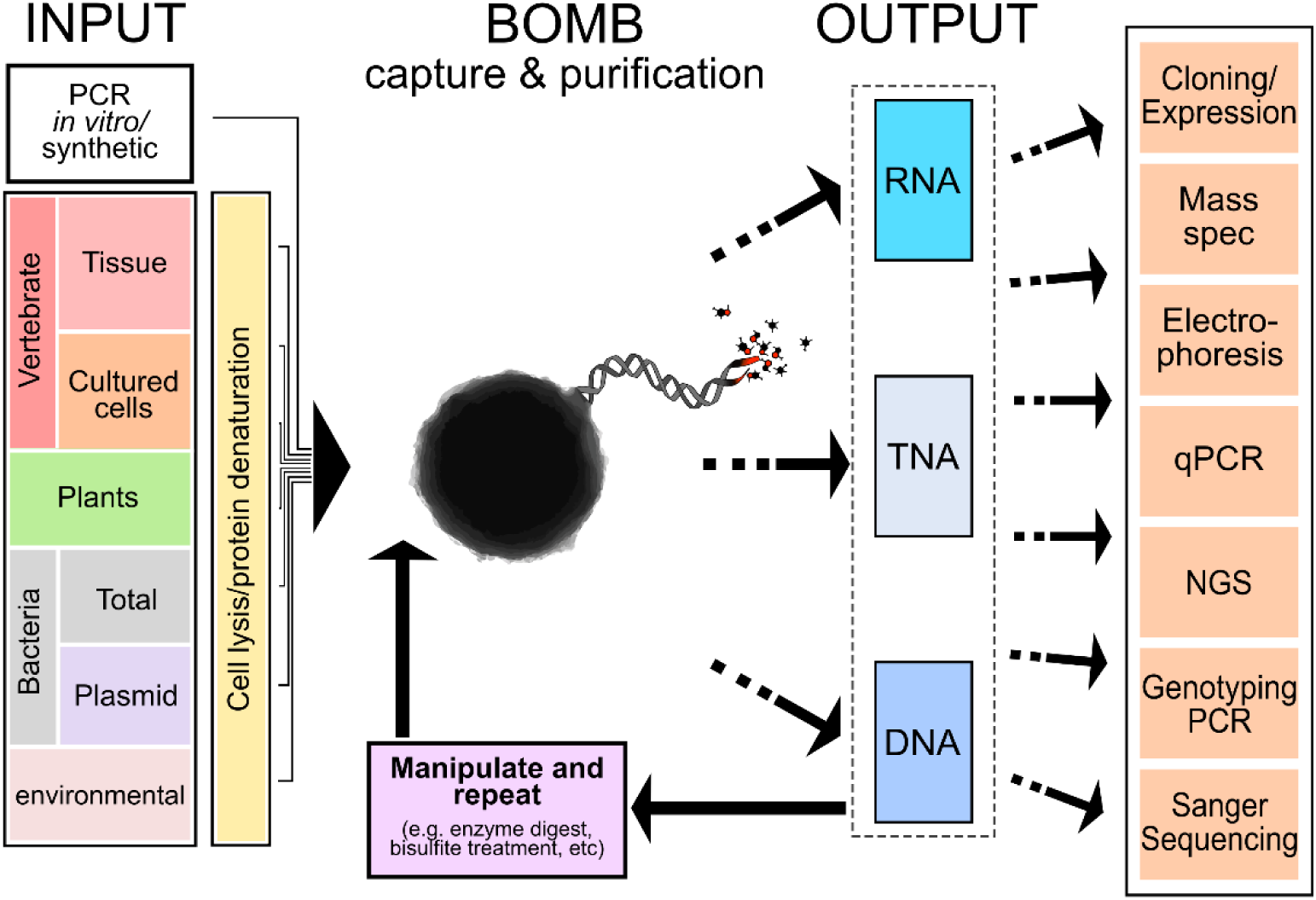
The BOMB platform is both modular and universal. Purification of nucleic acid from a wide variety of biological and synthetic input sources (left-hand-panel) can be performed using the BOMB platform. Following initial extraction using BOMB beads (central bead icon), the resulting nucleic acids, DNA, RNA or total nucleic acid (TNA) can be further manipulated and re-purified using BOMB protocols, and/or passed to final analysis methods or re-introduced back into a biological system.

### Protocol #4: Clean-up and size exclusion

The separation and purification of nucleic acids following enzymatic reactions is a necessary procedure in a variety of biochemical assays and was originally performed by precipitation with salts and alcohol [12–14]. However, these methods usually require long incubation steps of up to several hours for smaller molecules and, therefore, have been surpassed by rapid methods involving commercial silica columns or SPRI on magnetic beads [3]. A major advantage of SPRI bead methods is the ability to perform sequential enzymatic clean-ups in one tube in an efficient manner [15], thus greatly simplifying complex nucleic acid handling procedures such as DNA library construction for next-generation sequencing [4]. Moreover, because larger fragments precipitate to magnetic beads more efficiently than smaller ones in hydrophilic conditions, bead immobilisation can be used to select or exclude nucleic acids of particular sizes by varying the binding conditions.

Clean-up and size exclusion can be performed using self-synthesised BOMB beads, with either carboxyl-or silica-coating (Supplementary protocols #4.1 and 4.2), however, their optimal binding conditions differ. Whereas DNA is bound to carboxylated beads via molecular crowding with high concentrations of PEG-8000 and NaCl [16], binding DNA to silica beads utilises the altered affinity of the negatively charged DNA backbone to the silica surface in the presence of chaotropic salts [17,18]. We most commonly use silica-coated beads and guanidinium hydrochloride for capture. In doing so, we can selectively isolate fragments between 100 and 3000 bp (Fig 2A and S2A,B), depending on the amount of binding buffer used while recovering up to 95% of the input DNA (Fig S2C). Furthermore, utilising the 96-well format and the earlier described BOMB microplate racks, approximately 200 samples can be processed within 45 minutes by a single person. Lastly, we developed a separate bead protocol to purify DNA directly from agarose gels (Fig 2B, Supplementary protocol #4.3)

### Protocol #5 Bacterial plasmid extraction

Plasmid extraction from cultured *Escherichia coli* (*E. coli*) strains is probably one of the most common laboratory practices. In the late 1960s the first protocols for isolation of plasmid DNA were published [19–21], from which the alkaline lysis of bacterial cells in a slightly modified form became today’s primarily used method [17,22,23]. Numerous commercial kits are based on this technology, which employs either silica-packed columns or silica-coated magnetic beads. Both methods represent efficient and reliable techniques for DNA isolation with a reasonable cost of approximately 1.5 € per sample. However, for processing samples in a high-throughput scale the price can become a significant factor. Furthermore, the column-based protocols are not suited for processing high sample numbers as the capacity of common table centrifuges is usually limited to 24 tubes per run.

We have developed a high-throughput plasmid DNA isolation protocol using silica-coated magnetic beads. For this, bacterial colonies are grown in a 2.2 ml 96-well deep-well plate (Fig S3A), harvested and lysed with a modified alkaline lysis protocol. The plasmid DNA is then captured and immobilised with silica beads and remaining particles (cell debris, proteins, etc.) are washed out. Isolation of plasmid DNA (pUC19) with the optimised BOMB protocol (Supplementary protocol #5.1) yielded approximately 6 µg DNA per sample (Fig 2D,E). Amount, purity and quality of the extracted plasmid DNA are comparable to commercial preparations and the isolated DNA is suitable for both restriction digestion and Sanger sequencing (Fig S3B,C) or other common downstream applications. Using silica-coated BOMB beads we routinely process up to 200 samples in 3-4 hours for approximately 0.14 € per sample (Table 1).

**Table 1.** Cost comparison of BOMB protocols and commercial solutions for nucleic acid isolation and handling: Complete costs per 96 samples were calculated taking into account plastics, solvents and enzymes like DNase I. These costs were omitted for the kit content replacement cost column.

| Protocol | BOMB: Complete cost per 96 samples | BOMB: Kit content replacement per 96 samples | Price of commercial kits per 96 samples | Price advantage (kit/BOMB complete cost) |
| --- | --- | --- | --- | --- |
| Clean up | 5 € | 0.34 € | 155 – 218 € | 31 – 43x |
| Gel extraction | 11 € | 1.5 € | 155 – 228 € | 14 – 21x |
| Plasmid DNA extraction | 14 € | 5 € | 146 – 191 € | 10 – 14x |
| TNA extraction from cells | 11 € | 2.2 € | 349 – 1332 € | 32 – 121x |
| TNA extraction from tissues | 23 € | 6.5 € | 349 – 471 € | 15 – 20x |
| gDNA extraction | 12 € | 2.2 € | 183 – 354 € | 15 – 30x |
| Total RNA extraction | 32 € | 2.5 € | 440 – 592 € | 14 – 18x |
| Bisulfite conversion | 16 € | 5 € | 207 – 655 € | 13 – 41x |

### Protocols #6, 7 and 8: Purification of TNA, genomic DNA and total RNA

The isolation of TNA, genomic DNA or total RNA from bacteria and eukaryotic cells is a basic wet-lab technique and is the starting point for many molecular biology experiments. Most purification kits and techniques are designed and marketed to isolate either genomic DNA or total RNA. A classical method for DNA involves lysis of cells in a low-salt buffer with Proteinase K and a detergent, followed by phenol-chloroform extraction and ethanol precipitation [24]. For RNA, various protocols have been developed over the years [25–27], however, the most common involve guanidinium species as a strong denaturant that suppresses RNase activity [28] and facilitates binding of RNA to the silica bead. Many commercial kits are available for DNA and RNA as well, often utilising the addition of ethanol to bind the DNA or RNA to silica columns, with successive washes performed using a centrifuge. The commercial kits work well, however, the cost per sample exceeds 2 € and the flow-through column system is not easily adapted to 96-well format. Thus, experiments with a large number of samples are impossible for most laboratories.

We have found that for bacterial and mammalian cells without extensive extracellular protein, TNA can be efficiently purified using only a sarkosyl and guanidinium-isothiocyanate (GITC)-based buffer for protein denaturation and cellular lysis. (Fig 2C,F-H). For some cell-types such as sperm or spleen, addition of 2% β-mercaptoethanol may assist chromatin denaturation and inhibit RNA degradation, respectively. Following lysis, isopropanol is used to drive precipitation of the nucleic acid to the magnetic beads - BOMB silica beads work well for capture, as do carboxylated beads. The total volume is flexible, however the relative amount of each component should always be 2:3:4, that is, beads:lysate:isopropanol. Once the nucleic acid and beads are immobilised on a magnet, rapid isopropanol and 80% ethanol washes remove residual protein, GITC and other salts, allowing subsequent elution of the purified TNA. Enzymatic DNase or RNase treatment can be performed either before initial bead purification (e.g., during lysis steps, Fig 2F), or afterwards and subsequently re-purified on beads to produce solely RNA or DNA, respectively. Compared to commercial kits, this protocol performs well with a comparable yield of around 20 µg of genomic DNA isolated from 500K HEK293 cells (Fig 2G,H), while being slightly faster than most kits (∼1.5 h for 96 samples).

Cells derived directly from solid tissues and organs such as muscle and heart are not easily lysed in GITC without mechanical disruption. In this case, high-throughput lysis and protein digestion can be first performed in a low salt buffer using Proteinase K, followed by further denaturation in a more concentrated (1.5X) GITC buffer. While RNA is not well preserved using this high-throughput method, we have found that DNA purification works efficiently on a range of mammalian tissues (with either silica or carboxylated beads) compared to classical phenol-chloroform extraction (Fig 2L, S4). Further modifications to the initial lysis steps can be added in order to purify nucleic acid from a very broad range of sources. For example, with a small amount of prior mechanical disruption, TNA can be easily extracted from plants using just GITC lysis (Fig 2N,O, S6), or it can be combined with low-salt Proteinase K digestion. Lyticase pre-treatment can be employed for isolation of TNA from *Saccharomyces cerevisiae* cells (Fig 2M). It is even possible to extract good quality TNA from environmentally-derived sources, such as lake water (Fig 2P).

While TNA can be effectively purified using cells lysed in GITC buffer alone, some users may prefer the acid-guanidinium isothiocyanate-phenol solution [28] (Supplementary protocol #B or its commercial equivalent TRIzol®) for initial denaturation. Using silica-coated BOMB beads and TRIzol® solution we have isolated an average of 7.2 µg of pure total RNA from 500K HEK293 cells (Fig 2H,J) in 96 samples simultaneously, for a total cost of approximately 0.30 € per sample. The isolated RNA is intact and suitable for delicate analysis techniques like qPCR (Fig 2K) or RNAseq.

### Protocol #8: Bisulfite conversion

Methylation of DNA at the 5^th^ position of cytosines in the context of CG dinucleotides is probably the best studied epigenetic modification and likely plays a central role in defining vertebrate cellular identity [29–31]. Bisulfite sequencing is commonly used to study the distribution of 5-methylcytosine in genomes at single base resolution and it is considered the gold standard in the field [32]. This method is widely used and can be employed to sequence a wide variety of samples, ranging from single amplicons to whole genomes. It was developed in the 1990s by Frommer and colleagues [33] and uses sodium bisulfite to sulfonate unmodified cytosines at the 6^th^ position, followed by hydrolytic deamination and desulfonation in alkaline conditions, thereby converting unmodified cytosines to uracils. This reaction is far slower for methylated cytosines (or hydroxymethylated) at position 5, resulting in selective base conversion which can be detected by sequencing, allowing site-specific analysis of the methylation status by comparison to the original sequence (Fig S5A). One major drawback of this procedure is that bisulfite treatment causes DNA degradation, especially at high incubation temperatures. Therefore, over the years, the original protocol has been improved to accelerate the conversion procedure [34,35]. Multiple companies offer commercial kits for bisulfite conversion and some even use magnetic beads for high scalability, but they still cost more than 200 € per 96 sample plate.

The protocol we have developed follows a fast, optimised bisulfite conversion chemistry [34]; however, for the separation of converted DNA and following desulfonation steps, silica-coated BOMB beads are employed instead of column-based purification, thus allowing treatment of many samples in parallel. We have tested and optimised this protocol using 500-750 ng of DNA as the substrate (Fig S5B,C), but it has also worked well with input amounts ranging from 10 ng to 2 µg (data not shown). Sequencing of amplified converted DNA showed conversion rates comparable to commercial kits (∼99%). The whole procedure takes around 3-4 hours for 96 samples with a hands-on time of less than 1.5 h and costs less than 0.20 € per sample.

### Benefits and costs associated with the BOMB platform

Perhaps the most apparent benefit of the BOMB system is economic. Commercial column-based nucleic acid extraction kits are commonly used in laboratories. Depending on sample type, the nature of the nucleic acid to be purified and the vendor, the price per sample ranges between 1-14 € (Table 1). A single synthesis of silica BOMB beads is sufficient for >40,000 extractions of RNA, genomic DNA or plasmid DNA, bringing the costs of nucleic acid extractions to less than 0.05 € (clean-up) to 0.32 € (total RNA extraction) per sample. These costs have been calculated using high-yield extractions based upon deep-well plates (1-2 ml); however, further significant cost savings can be made by scaling reaction volumes down to 0.2 ml PCR tubes and plates. Being at least 10-20 times cheaper than commercial column-based protocols, BOMB methods are suitable for large scale experiments on a budget. Generally, the most expensive aspects of the BOMB platform are enzymes, like DNase I or RNase A; however, these costs can be greatly reduced, if purification of a single nucleic acid species is not required. For example, routine PCR and even bisulfite sequencing can be performed on TNA without the need to remove RNA. Other significant costs associated with the BOMB platform include washing solvents such as ethanol and disposable tips and plates, however, these are usually not included in commercial kits and have to be supplied by the user. Most of the necessary chemicals required for creating the BOMB platform are readily available in standard molecular biology laboratories, and the rest can be purchased from a range of vendors. For the sake of rigour, our protocols list the suppliers we have used, however, we expect that good quality chemicals from an alternative source will show similar performance.

We have developed BOMB protocols as a consequence of an immediate need to process hundreds of samples on a tight budget and with limited manpower. By using magnets to immobilise nucleic acid captured on magnetic beads, centrifugation steps are eliminated. As such, these methods are highly scalable and allow easy processing of multiple batches of 96 samples in parallel by a single researcher with a multi-channel pipette. Because of the 96-well format and the lack of dependence on a centrifuge, these protocols are also automation friendly and can be adapted to liquid handling robot systems if available. Compared to kits with single spin columns, our protocols are much faster when processing many samples simultaneously. However, even when processing only 1-10 samples, most of our protocols are at least as fast as commercial alternatives while retaining the cost advantage, high yield and quality of the isolated nucleic acids.

### Switching to a bead-based lab

We have found that the time invested in switching to bead-based protocols is very quickly returned and changing over to a primarily bead-based laboratory actually simplifies and accelerates many experimental processes. For example, a researcher may wish to undertake genome-wide methylation and expression analysis from the same cultured cell source. Here, a single TNA extraction can be performed - half of the material can be used directly for bisulfite conversion, while the rest can be DNase-treated and further purified to obtain pure RNA, which can then be processed into RNA-sequencing libraries.

Converting to a low-cost bead-based system represents an opportunity to transform the research culture of any laboratory in a profound way. By having the capacity to do high-throughput experiments without extra cost, researchers may gain the ability to study entire populations instead of merely sampling, include extra control samples they otherwise would have to forego, or consider analysing separate cell populations rather than relying on averages generated from bulk tissue. Integration of next-generation sequencing with the BOMB platform provides additional synergy in these respects. Double-ended indexing allows pooling of potentially thousands of individual samples and amplicons in a single run, yet, most laboratories are unable to capitalise on this empowering aspect because, until now, they were not equipped to process samples of this number.

A striking feature of the BOMB platform is its versatility and universality (Fig 3). Nucleic acid can be purified from a diverse set of biological and non-biological sources and then passed on to any number of additional BOMB-based protocols, ranging from genotyping to bisulfite sequencing, library construction and clean-up/size selection following PCR. Each step of these modular pipelines magnifies the benefits of the BOMB platform - we recently purified human cell line RNA, constructed cDNA, then amplified and cloned >100 human genes within 3-weeks using BOMB protocols (Oberacker et al., in preparation). Because multiple rounds of plasmid DNA isolation, screening for positive clones, and sub-cloning was performed, the additive advantage of using BOMB protocols in a 96-well format over conventional targeted cloning approaches was immense.

Initially, all that is required for a switch to the BOMB platform are beads and a magnetic rack, so getting started is easy. And while we are confident all laboratories can make their own beads, one barrier to starting with the BOMB platform can be removed by buying commercially available silica-or carboxylate- functionalised beads. To ease the transition to a BOMB-based lab we have included a forum section on our website (https://bomb.bio/) where researchers can ask questions and get help from other users, serving as a platform for further development of protocols particularly for unique species and applications. Furthermore, the community can use this platform to expand the repertoire of procedures and sample types in a collaborative manner by discussing existing protocols, providing adaptations and completely new protocols.

### Outlook

Here, we provide a set of simple, step-by-step protocols for extraction, purification and manipulation of nucleic acids from various sources. These protocols can serve as a starting platform for further development of other functionalised magnetic nanoparticles, as well as protocols tailored to the specific experimental needs of the users. Currently our focus is on nucleic acids, however, we expect further bead-based protocols will continue to be developed for more diverse applications. For example, both the carboxylate and silica coatings can be further chemically derivatised by attaching additional functional groups, like cofactors, proteins or antibodies (for example coating with protein G), which will in turn allow additional functionalities of the beads. Our community focussed website and forum will facilitate this development, allow troubleshooting, reagent sharing and the distribution of new user-developed protocols. We envision that better access to magnetic bead technology will drive greater efficiency of research in the life sciences, and further empower our collective quest for knowledge.

## Acknowledgements

The authors would like to thank all members of the Jurkowski and Hore laboratories for helping to optimise and test the BOMB protocols. We are also indebted to Ken Wyber (Otago Polytechnic) for help with laser cutting magnetic plates. We are grateful to Dr. Renata Jurkowska for critical reading of the manuscript. We would like to thank the wider research community for offering unpublished information and resources concerning magnetic bead preparation and utility, in particular, Dr Ethan Ford, Dr James Hadfield, Dr Brant Faircloth, Dr Nadin Rohland and associated authors.

## Author’s contribution

The idea was conceived by TPJ and TH. Protocol setup and optimisation was led by PO, PS, DB, TPJ and TH, with contributions from SH, JF, VM, LS, VJS, G-JJ and FvM. Laser cutting designs were contributed by SRH. The electron microscope analysis was done by KH. The website and its content were created by TM, PS, PO, TPJ and TH. The manuscript was written by TPJ, TH, PS and PO. All authors contributed to the editing of the manuscript and approved its final version.

## Supporting Information

**Fig S1:**
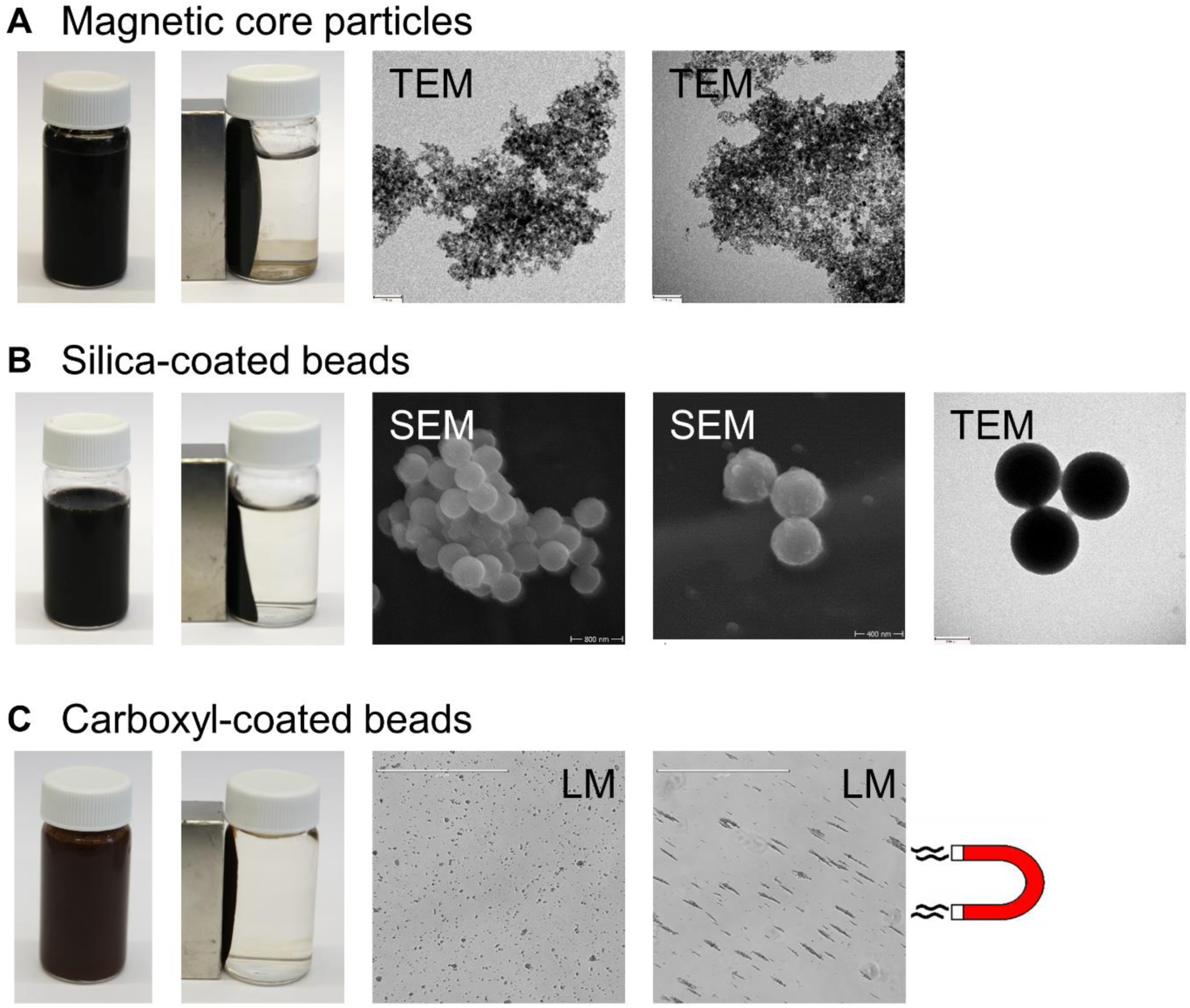
BOMB magnetic beads. (A) Magnetic core particles (see protocol #1) in transmission electron microscopy (TEM). (B) Silica-coated magnetic beads (see protocol #2.1) in scanning electron microscopy (SEM) and TEM. (C) Carboxyl-coated magnetic beads (see protocol #3.1) in light microscopy (LM), with and without an applied magnet.

**Fig S2:**
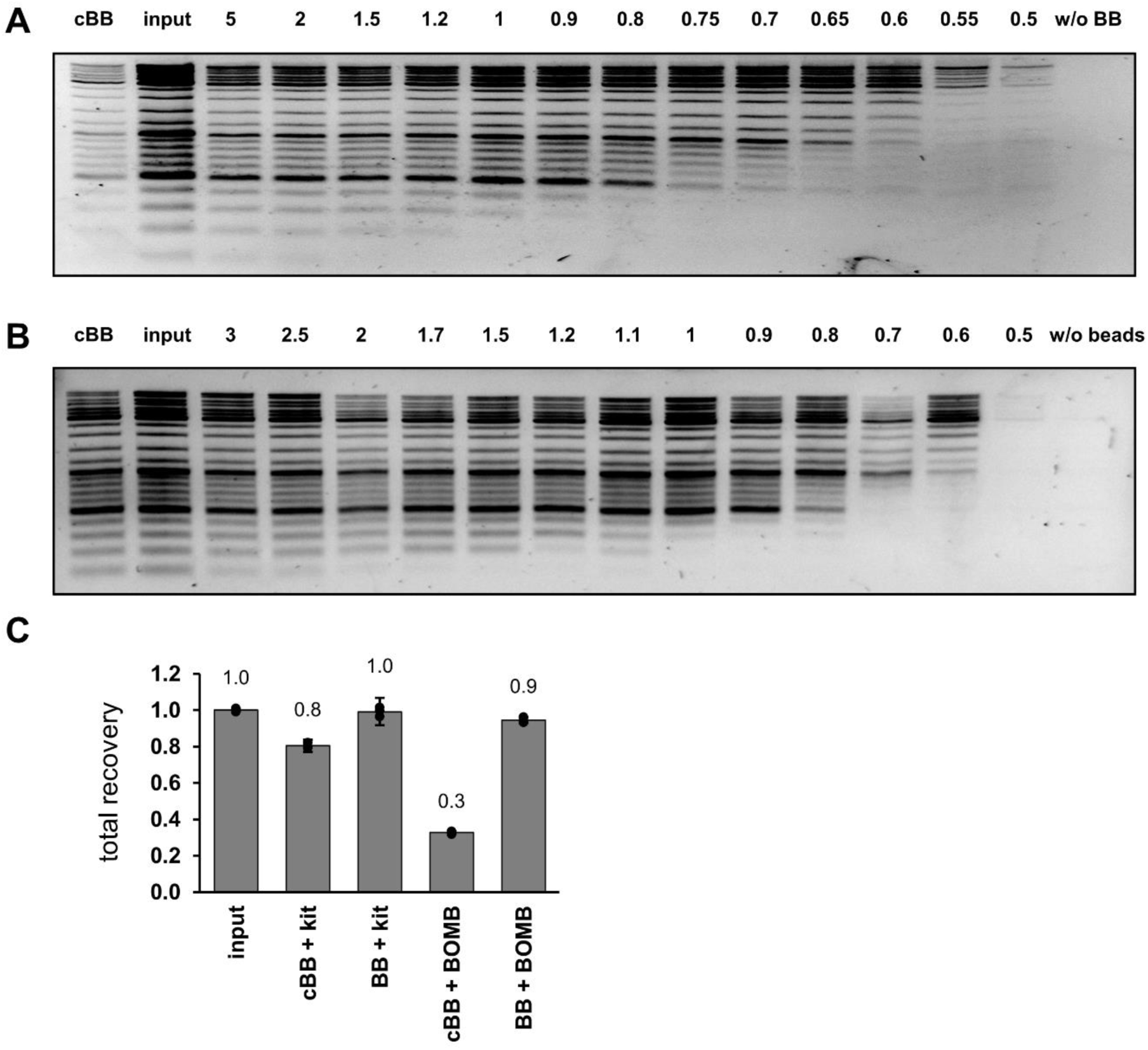
Clean-up and size exclusion of DNA. (A) Size exclusion of GeneRuler DNA Ladder Mix (Thermo) using silica-coated magnetic beads (see protocol #4.1). Different volumes of binding buffer (BB) compared to sample volume were used to achieve size exclusion; as a comparison 2 volumes of commercial BB (cBB) were used, or no binding buffer was included (w/o BB) (B) Size exclusion of GeneRuler DNA Ladder Mix using carboxyl-coated magnetic beads (see protocol #4.2). Different volumes of binding buffer (BB) compared to sample volume were used to achieve size exclusion; as a comparison 3 volumes of commercial BB (cBB) were used, or no beads were included (w/o beads). (C) Total recovery of ∼6 µg plasmid DNA (input) using either a commercial kit that includes silica-packed columns (kit) or the #4.1 clean-up protocol with silica-coated beads (BOMB). For binding, either commercial binding buffer (cBB) or the binding buffer (BB) described in the BOMB protocol above was used. Error bars represent standard deviation, n=3.

**Fig S3:**
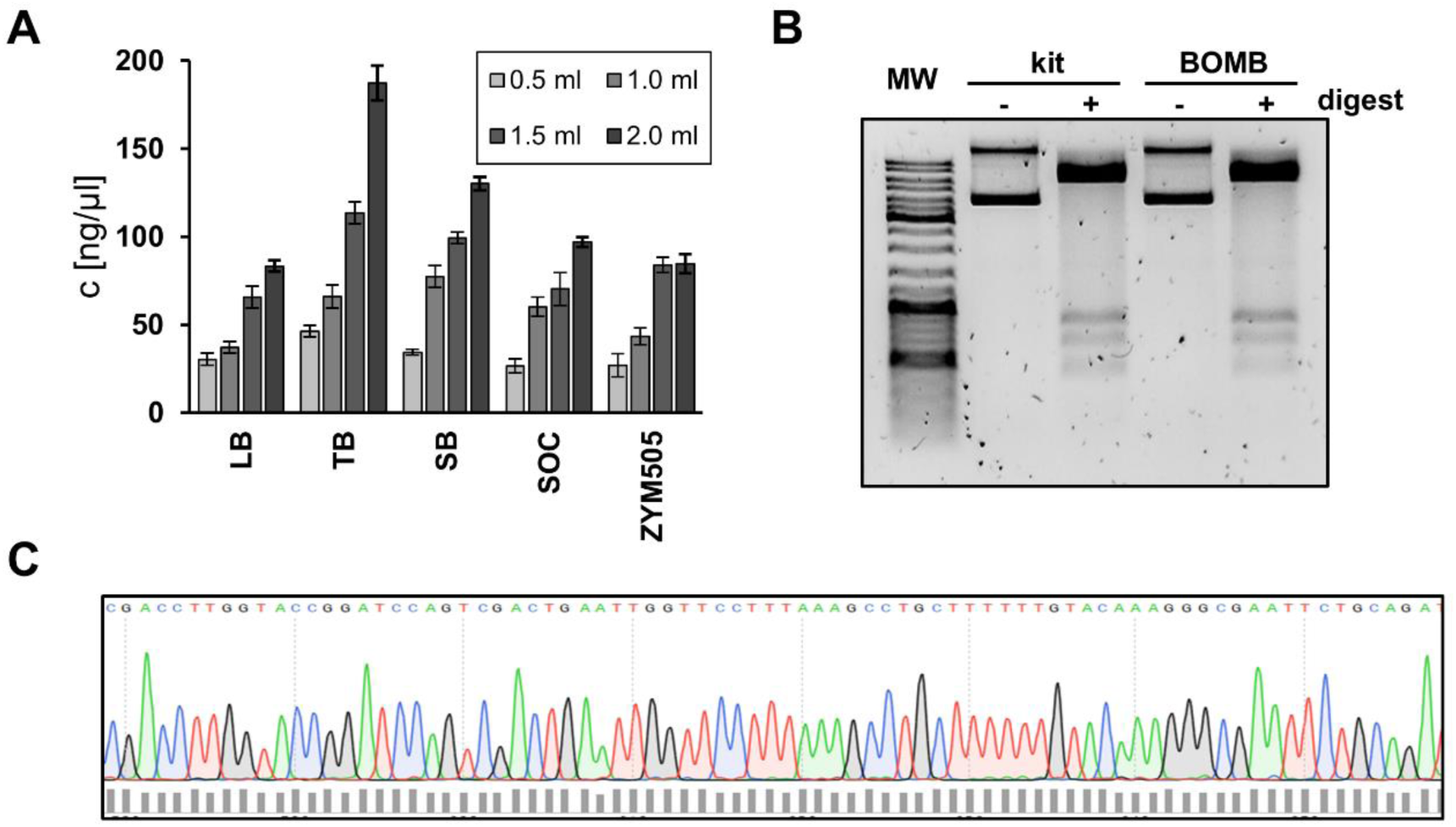
*Optimisation and quality control of #5.1 BOMB plasmid extraction from* E. coli. (A) Optimisation of reaction volume and media. One colony was picked and used to inoculate a 5 ml pre-culture containing LB media and ampicillin. The culture was incubated at 37 °C and 250 rpm until an OD of 0.6 was reached. 5 µl of the pre-culture was used to inoculate different volumes (0.5 ml, 1.0 ml, 1.5 ml and 2.0 ml) of a variety of growth medias (LB, TB, SB, SOC and ZYM505; including the respective antibiotic) in a 2.2 ml 96-well culture plate (Sarstedt). The plate was sealed with the lid of a 6-well cell culture plate, so a decent exchange of air was possible, and incubated at 37 °C and 250 rpm for 22 h. Plasmid DNA was than isolated with the BOMB plasmid extraction protocol and the resulting concentration (c [ng/µl]) was determined. Error bars represent the standard deviation, n = 3. (B) Comparison of commercially purified plasmid DNA (kit) and BOMB extracted DNA (BOMB) with (+) and without (-) restriction enzyme digestion. MW: GeneRuler DNA Ladder Mix (Thermo Scientific). (C) Exemplary sequencing trace of a BOMB extracted plasmid via Sanger sequencing. A sequencing read length of at least 800-1000 bp is typically observed.

**Fig S4:**
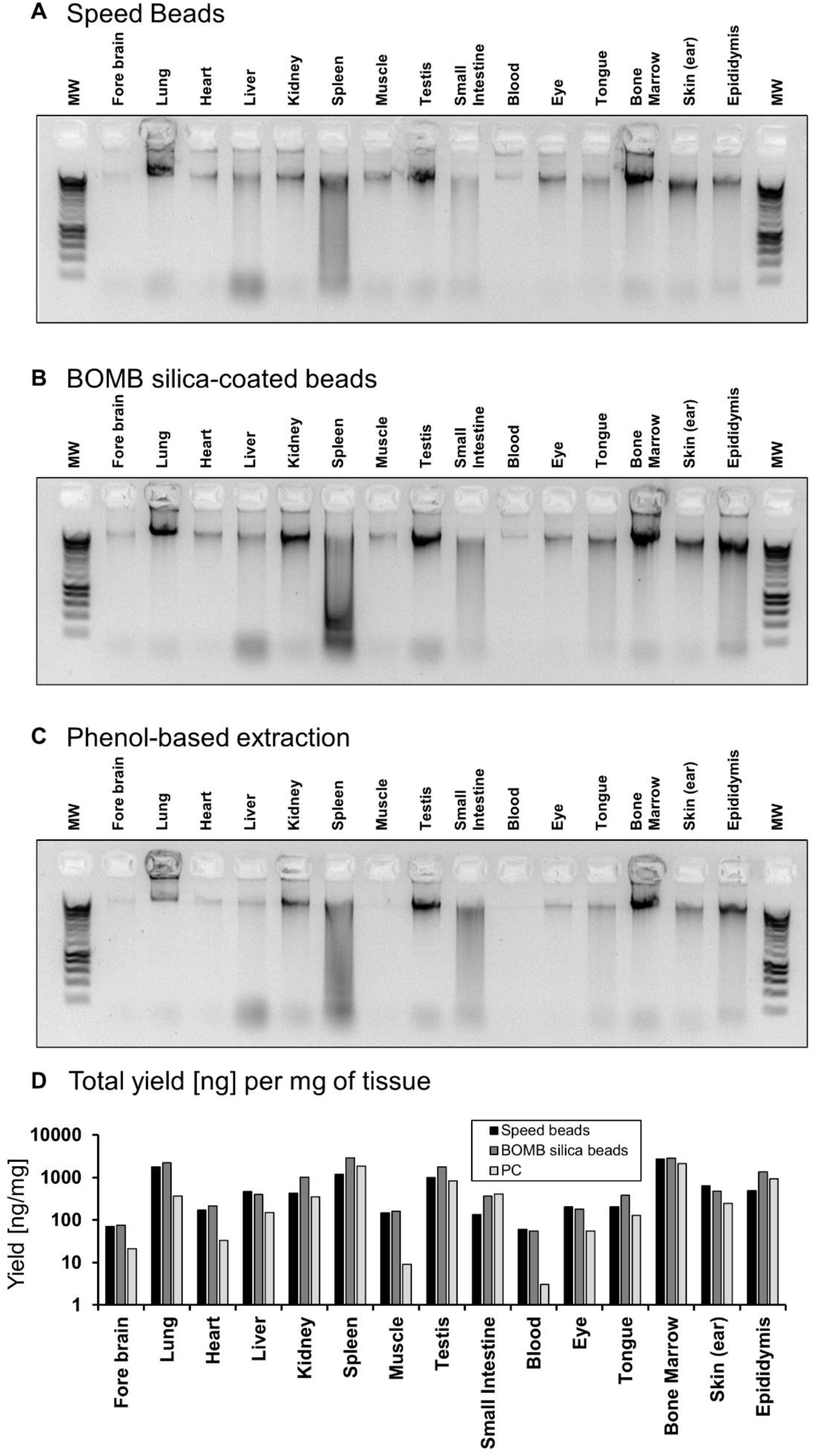
Genomic DNA isolation from various rabbit tissues. Genomic DNA was isolated from the indicated tissues of a 12-hr deceased rabbit, using protocol #6.3 and (A) Speed Beads or (B) BOMB silica beads. A comparison to (C) Phenol-chloroform based extraction is also shown. MW in all panels represents Hyperladder I (Bioline). Inevitably, some tissues (like bone marrow) produce far greater (D) yields per mg of input material, compared to other tissues. However, the bead-based methods generally outperform phenol-chloroform extractions in our hands. Note, rabbit tissues were not preserved immediately after animal death, hence why tissues like spleen have experienced some DNA degradation.

**Fig S5:**
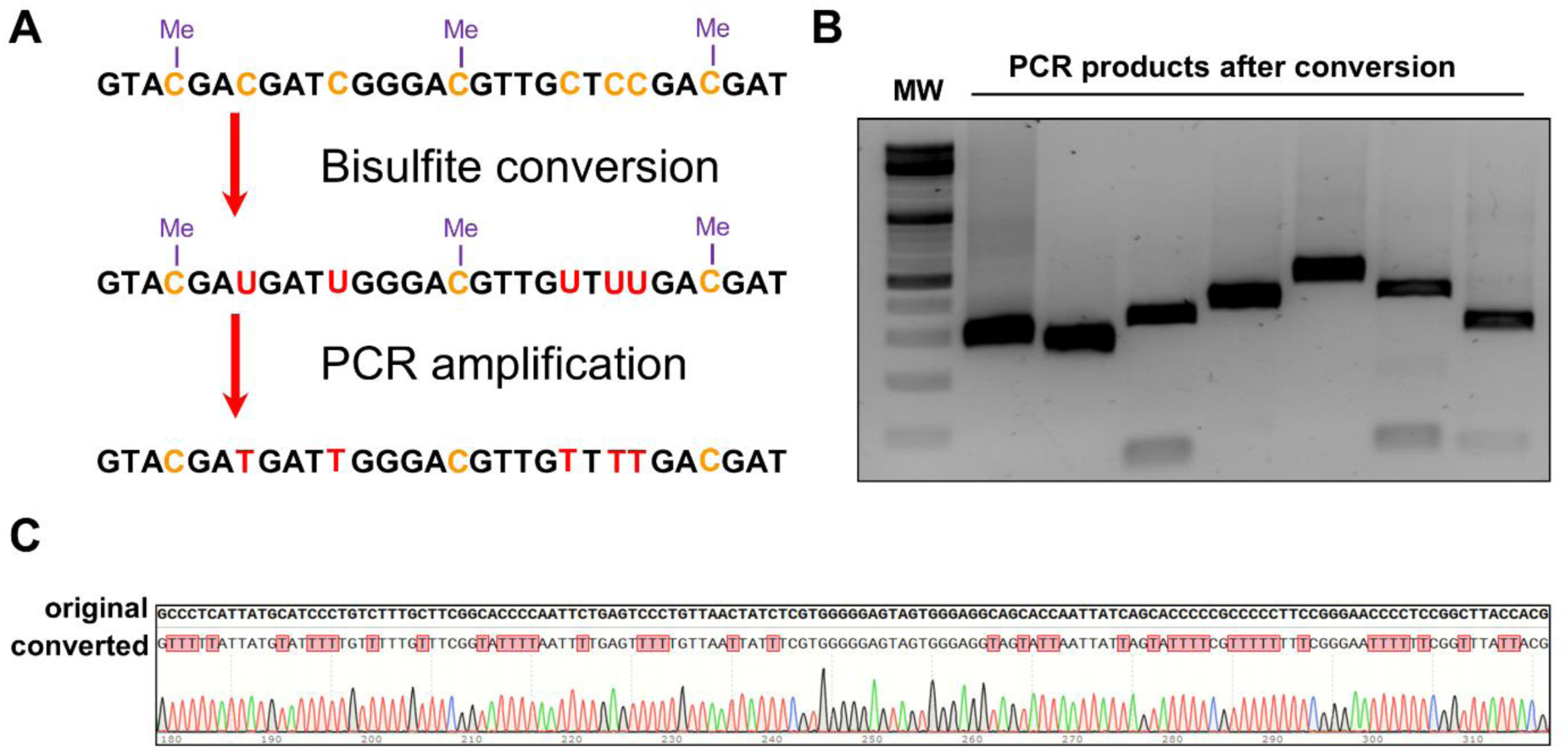
Bisulfite conversion. (A) Scheme of DNA methylation analysis using bisulfite conversion. (B) Agarose gel after PCR amplification of bisulfite converted human DNA. Multiple primer pairs with expected product sizes between 221 and 435 bp were tested successfully. MW: GeneRuler DNA Ladder Mix (Thermo Scientific). (C) Sequencing trace of a PCR amplified bisulfite converted sample, aligned with the original, unconverted sequence. All non-CG cytosines were successfully converted. Conversion rate is ∼99% as measured by Sanger sequencing after PCR amplification.

**Fig S6:**
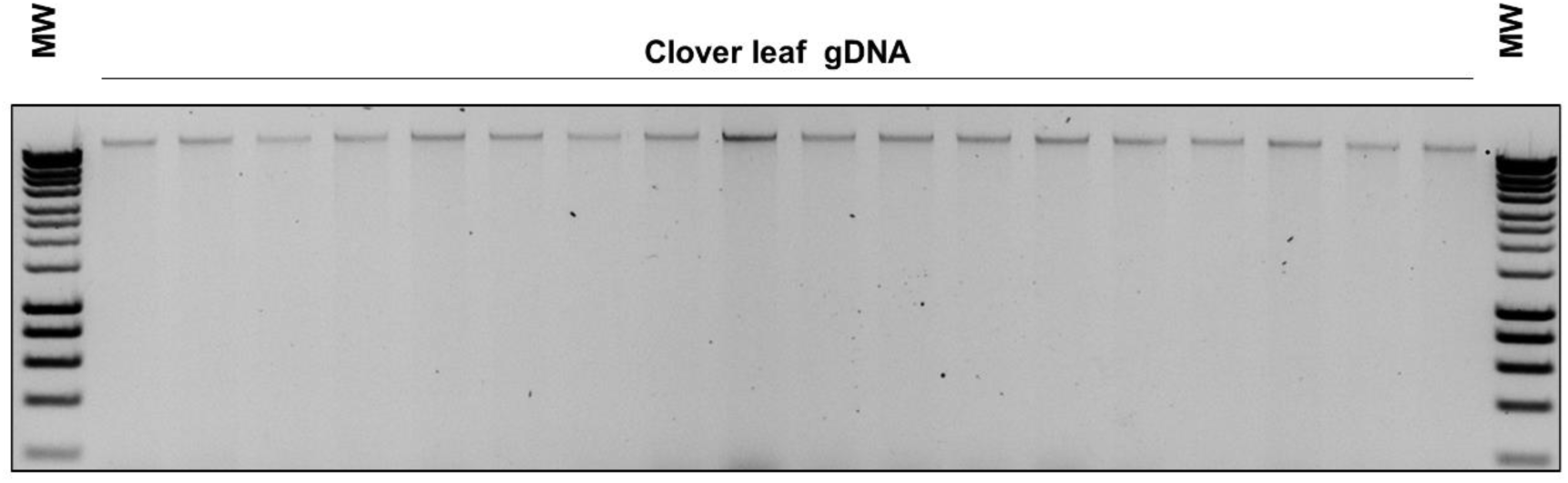
*Example genomic DNA isolation from clover leaves (*Trifolium repens). Genomic DNA was isolated from individual leaves (∼ 5mg of tissue) using protocol #6.4 (high-throughput). A subset of DNA samples (18) from 96 extractions, represented in Figure 2O, were run on an agarose gel. MW: Hyperladder I (Bioline).

## Abbreviations

BOMB: Bio-On-Magnetic-Beads
SPRI: Solid-Phase Reversible Immobilisation
MNP: magnetic nanoparticle

